# Regulation of multiple paralogs of a small subunit ribosomal protein in *Francisella tularensis*

**DOI:** 10.1101/2025.06.29.662229

**Authors:** Sierra S. Schmidt, Alexandra R. Farah, Aisling Macaraeg, Daniel Floyd, Hannah S. Trautmann, Kathryn M. Ramsey

## Abstract

*Francisella tularensis* is a highly infectious human pathogen that must replicate inside macrophage to cause disease. The ribosomes of *F. tularensis* can incorporate one of three different paralogs for the small ribosomal subunit protein bS21. One of these paralogs positively impacts translation of key virulence genes and promotes intramacrophage replication. Although ribosomal bS21 content influences *F. tularensis* virulence, the factors that control bS21 paralog production are not well understood. Here, we reveal that all three bS21 proteins influence the transcript abundance of the paralog important for virulence, bS21-2. In contrast, the other bS21 paralogs (bS21-1 and bS21-3) do not affect their own production. We further determined that the leader sequence of the bS21-2 mRNA is sufficient for bS21-mediated repression of mRNA abundance, suggesting that bS21-2 is autogenously regulated. Yet we determined that the increase in bS21-2-encoding mRNA is not reflected by increased protein production, suggesting that translation of this transcript is controlled by other factors. Finally, we found that bS21-2 exerts at least some of its effects on the bS21-2 transcript by decreasing its stability. Together, our findings suggest that *F. tularensis* integrates multiple signals into a regulatory network to control the appropriate production of each bS21 paralog, and particularly the paralog important for virulence, bS21-2. This regulatory network in turn may control ribosomal heterogeneity and virulence gene expression.

## Introduction

Among the cellular processes, translation is the most energetically costly (Hu et al., 2020). Part of this cost is the assembly of the large molecular machine that catalyzes protein synthesis, the ribosome. Accordingly, cells have complex regulatory networks to balance the production of rRNA and ribosomal proteins (r-proteins). For example, operons encoding multiple r-proteins are often regulated by translational feedback, a form of autogenous gene regulation. In these networks, if there is an overabundance of one of the first r-proteins to bind rRNA in ribosome assembly (primary binding r-proteins), this excess protein may bind and inhibit translation of their own mRNA, sometimes via an interaction with a structure analogous to its rRNA binding site (Meyer, 2018; Nomura et al., 1984).

It is common for bacteria to encode multiple paralogs for at least one ribosomal protein, which can lead to heterogeneity in ribosome composition (Yutin et al., 2012). This raises the question of how production of r-protein paralogs is coordinated and balanced. Significant advances have been made in understanding this coordination in the case of zinc-regulated r-protein paralogs. Diverse bacteria encode multiple paralogs for one or more r-proteins, one of which is capable of coordinating zinc and one (or more) others which does not (Makarova et al., 2001). The paralog that coordinates zinc is produced in zinc-replete conditions but in lower-zinc environments, zinc-sensitive transcription factors allow their production and these alternate paralogs are subsequently incorporated into ribosomes (Graham et al., 2009; Hemm et al., 2010; Prisic et al., 2015; Ueta et al., 2020). This mechanism of paralog switching, dependent on the concentration of a divalent cation and the de-repression of a transcription factor, is relatively straightforward and is thought to allow dynamic control of intracellular zinc concentrations (Cheng-Guang and Gualerzi, 2021; Nanamiya et al., 2004; Panina et al., 2003; Prisic et al., 2015; Shin and Helmann, 2016). However, it is not understood how production of r-protein paralogs that are unrelated to zinc coordination are regulated.

The small ribosomal subunit protein bS21 (previously S21) is small (∼8 kDa) and does not encode the CxxC domain required for zinc coordination (Ban et al., 2014). It is one of the last proteins assembled and can be easily exchanged (Held et al., 1973; Pichon et al., 1975; Robertson et al., 1977). Located next to uS11 and bS18, it creates part of the small subunit platform domain and forms part of the mRNA exit channel, directly adjacent to mRNA leader sequences during initiation. Many bacteria do not encode a bS21 homolog, indicating that it is dispensable for protein synthesis (Galperin et al., 2021). But multiple studies reveal that it plays a role in translation initiation, with regulatory effects on initiation most clearly defined in *Flavobacterium johnsoniae* (Chang and Craven, 1977; Duin and Wijnands, 1981; Jha et al., 2020; McNutt et al., 2023; Trautmann et al., 2023; Trautmann and Ramsey, 2022). The current structural and functional studies raise the possibility that bS21 acts not as a core component of the ribosome but instead as a regulator of translation initiation.

The highly infectious intracellular human pathogen *Francisella tularensis* encodes three distinct paralogs of the small ribosomal subunit protein bS21, none of which are predicted to coordinate zinc or other divalent cations. We have demonstrated that *F. tularensis* ribosomes are heterogenous with respect to bS21 content and that changes in bS21 content impact gene expression, dependent on specific mRNA leader sequences (Trautmann et al., 2023; Trautmann and Ramsey, 2022). Additionally, ribosomes with one paralog, bS21-2, are uniquely important for both production of a critical virulence factor (the type VI secretion system) and intramacrophage survival, which is essential for *F. tularensis* to cause disease (Trautmann and Ramsey, 2022).

How *F. tularensis* cells balance production of the three zinc-free bS21 paralogs is not clear. bS21 is not a primary binding r-protein (it is among the last to be assembled into the ribosome) and none of the *F. tularensis* bS21 paralogs are encoded in operons with other primary binding r-proteins (Mizushima and Nomura, 1970; Trautmann and Ramsey, 2022). However, there is evidence that in *F. tularensis*, bS21-2 regulates its own production; in cells without bS21-2 there is a large increase in mRNA corresponding to the leader sequence for the bS21-2 gene, *rpsU2*, and the gene immediately downstream, *yqeY*, which can be complemented by ectopic expression of bS21-2 (Trautmann and Ramsey, 2022).

There is limited information regarding control of bS21 in other organisms. *Escherichia coli* encodes only a single bS21 protein which is not reported to be subject to translational feedback regulation (Nomura et al., 1984; Takata, 1978). In *F. johnsoniae*, the single bS21 protein regulates its own production, but through a mechanism that appears specific to certain groups in the Bacteroidia phylum. Specifically, the C-terminal region of *F. johnsoniae* bS21 is conserved among Bacteroidia species and contributes to the sequestration of the anti-Shine-Dalgarno (ASD) (Jha et al., 2020; McNutt et al., 2023). This sequestration precludes ribosomes from being responsive to Shine-Dalgarno (SD) sequences, but the transcript encoding bS21 is the only *F. johnsoniae* mRNA with a strong SD. Thus, when bS21 is depleted and the ASD becomes accessible, translation of the bS21-encoding transcript increases (McNutt et al., 2023). However, this mechanism for self-regulation appears limited to the bS21 paralogs in Bacteroidia with similar conserved C-terminal sequences and structures.

We initiated our studies of bS21 regulation in the live vaccine strain (LVS) of *F. tularensis* by examining control of bS21-2. When comparing RNA abundance in cells with and without bS21-2, we found that the presence of any of the three *F. tularensis* bS21 paralogs reduces steady-state levels of the bS21-2-encoding transcript, *rpsU2*. This regulation appears to be unique to *rpsU2*, as we did not find that bS21-1 or bS21-3 similarly control their own production. Using reporter assays, we found that despite the increase in *rpsU2* transcript in cells without bS21-2, there is no concordant increase in protein abundance, suggesting that other regulatory mechanisms control *rpsU2* translation. Additional reporter assays demonstrated that while the 5’ UTR of *rpsU2* is sufficient for control of transcript abundance by bS21-2, the first 38 nucleotides (nt) of the *rpsU2* transcript, predicted to encode two stem-loops, are dispensable for regulation. Finally, we found that the presence of bS21-2 leads to decreased stability of the *rpsU2* transcript, which may be sufficient to explain the impact of bS21-2 on steady-state abundance of the *rpsU2* transcript. Our results suggest that control of the bS21 paralogs in *F. tularensis*, and particularly of bS21-2, is complex and regulated by multiple factors.

## Results

### bS21 paralogs control abundance of the bS21-2-encoding mRNA

Previous studies used RNA-Seq to examine the effects of bS21-2 deletion on mRNA abundance (Trautmann and Ramsey, 2022). Notably, the gene with the largest change in transcript abundance, *yqeY*, is encoded in the same operon and immediately downstream of *rpsU2*, the bS21-2 gene. Loss of bS21-2 leads to a 6-fold increase in *yqeY* transcript, which can be complemented by ectopic expression of bS21-2. Further analysis reveals that the region corresponding to the 5’ UTR of *rpsU2* is increased 9-fold in the absence of bS21-2 (**Fig 1**). These results indicate that bS21-2 functions as a negative regulator of the *rpsU2* operon.

**Figure 1.**
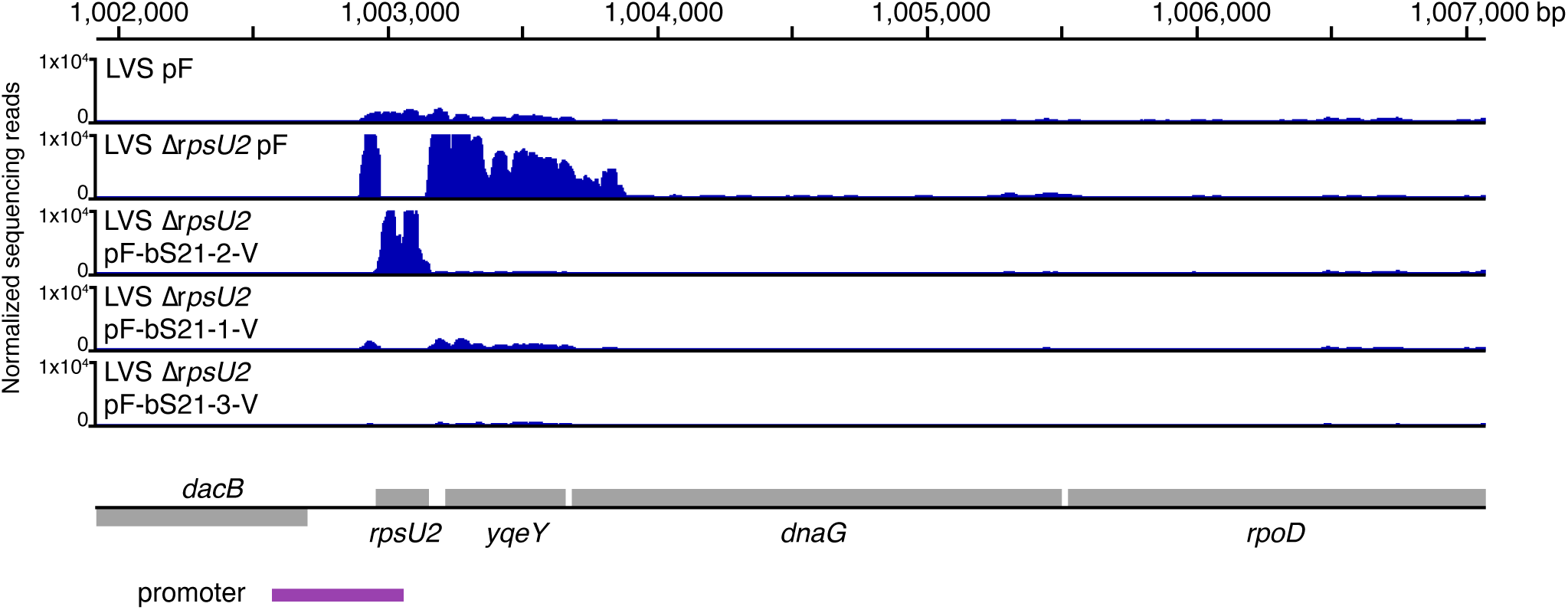
All three *F. tularensis* bS21 paralogs negatively regulate the *rpsU2* operon. Normalized transcript abundance reads from RNA-Seq experiments in the area surrounding the *rpsU2* operon. Data from cells with bS21-2 (LVS pF), cells lacking bS21-2 (LVS Δ*rpsU2* pF), and cells lacking bS21-2 but ectopically expressing either bS21-2, bS21-1, or bS21-3 (LVS Δ *rpsU2* pF-bS21-2-V, pF-bS21-1-V, pF-bS21-3-V, respectively). Y-axis is truncated at 1×10^4^ for clarity. Grey rectangles represent genes and those above the line indicate genes encoded on the positive strand; those below the black line represent those encoded on the negative strand. The purple box indicates an experimentally-determined promoter region (Ramsey et al., 2015).

Additional RNA-Seq data revealed that the other bS21 paralogs can also reduce the amount of *rpsU2* operon transcript. In particular, ectopic expression of either bS21-1 or bS21-3 in cells lacking bS21-2 results in reduction of the *rpsU2* operon transcript to levels similar to those found in wild-type cells (**Fig 1**). Thus, all three bS21 paralogs function similarly with respect to reducing the abundance of mRNA corresponding to the *rpsU2* operon.

### bS21-1 and bS21-3 do not control their own production

While loss of bS21-2 did not affect the abundance of the transcripts encoding bS21-1 or bS21-3 (*rpsU1* and *rpsU3*, respectively; (Trautmann and Ramsey, 2022), finding that bS21-2 regulates its own mRNA raises the possibility that other bS21 paralogs in *F. tularensis* may also regulate their own transcripts. To test this possibility, qRT-PCR was used to assess the relative amount of transcripts encoding these proteins in cells with and without the r-protein of interest. Specifically, we compared transcript abundance of the 5’ UTR of *rpsU1* in cells with and without bS21-1 (**Fig 2A**) and the 5’ UTR of *rpsU3* in cells with and without bS21-3 (**Fig 2B**). In contrast to our findings regarding the control of bS21-2, loss of the other two bS21 paralogs does not significantly impact the abundance of their own transcripts.

**Figure 2.**
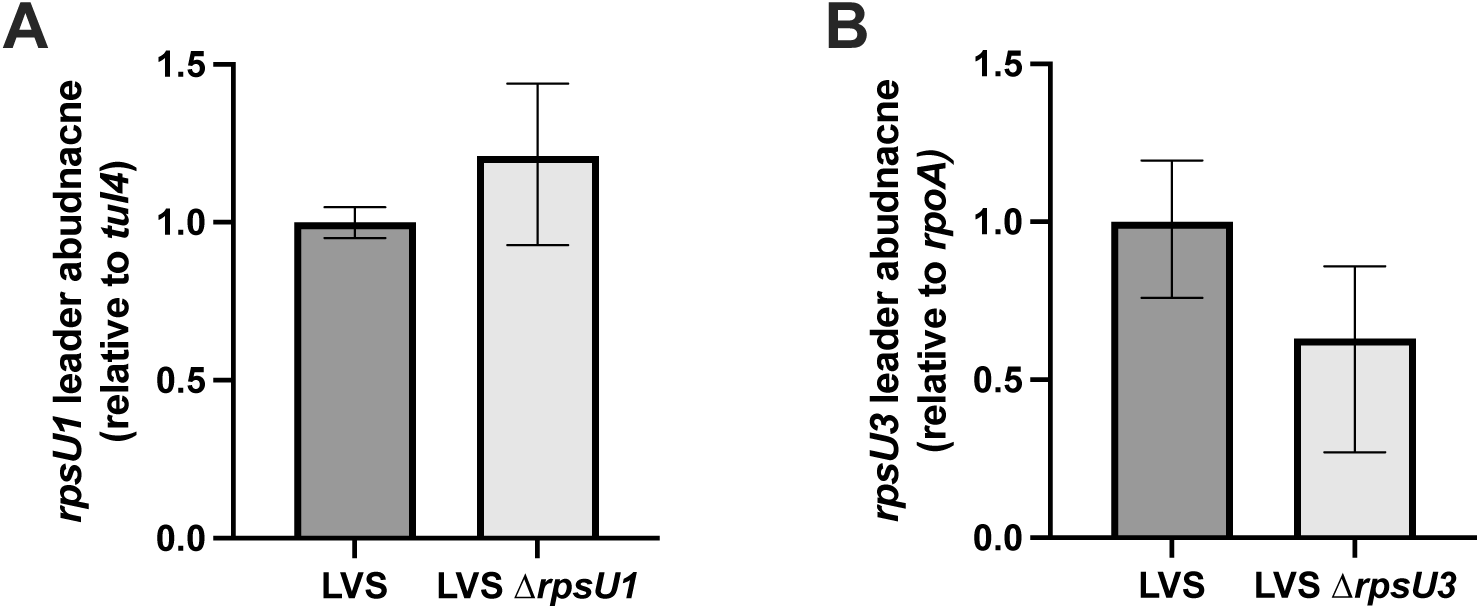
bS21-1 and bS21-3 do not regulate their own transcript abundance. (A) Transcript abundance of the *rpsU1* 5’ UTR in either wild-type cells or cells lacking *rpsU1*, relative to *tul4* (a control gene). (B) Transcript abundance of the *rpsU3* 5’ UTR in either wild-type cells or cells lacking *rpsU3*, relative to *rpoA* (a control gene with lower transcript abundance than *tul4*). (A and B) Differences did not reach statistical significance by t test. Experiments were repeated at least twice in biological triplicate and data from a representative experiment are shown.

### Transcript abundance and translation of the *rpsU2* operon are regulated independently

We constructed several reporter fusions to examine what sequence elements lead to regulation of bS21-2 production. In particular, we compared reporter activity from a fusion of the *tul4* promoter and 5’ UTR (a gene with expression unaffected by bS21-2; Trautmann and Ramsey 2022) to *lacZ* or a similar fusion with the *rpsU2* promoter and 5’ UTR in cells with or without bS21-2 (**Fig. 3A**). These translational reporters, which include DNA specifying the first six codons of the respective genes, were incorporated in single copy into the chromosome at the *att*Tn7 site. We subsequently measured abundance of the *lacZ* transcript by qRT-PCR and ß-galactosidase abundance using ß-galactosidase assays. In cells with the control (*tul4*) reporter, the amount of *lacZ* transcript is essentially the same, regardless of bS21-2 presence (**Fig 3B**). But in cells lacking bS21-2, the *lacZ* transcript produced from the *rpsU2* reporter increases by ∼9-fold (**Fig 3B**). This provides evidence that our translational fusions recapitulate control of the native *rpsU2* operon by bS21-2. However, translation of the *lacZ* mRNA with the *rpsU2* leader sequence was not affected by bS21-2 in the same way. Despite the ∼9-fold increase in *lacZ* transcript, there was only ∼30% more ß-galactosidase activity in cells without bS21-2 compared to cells with bS21-2 (**Fig 3B**). Thus, we determined that relative changes in bS21-2-controlled transcript abundance are not reflected in translation.

**Figure 3.**
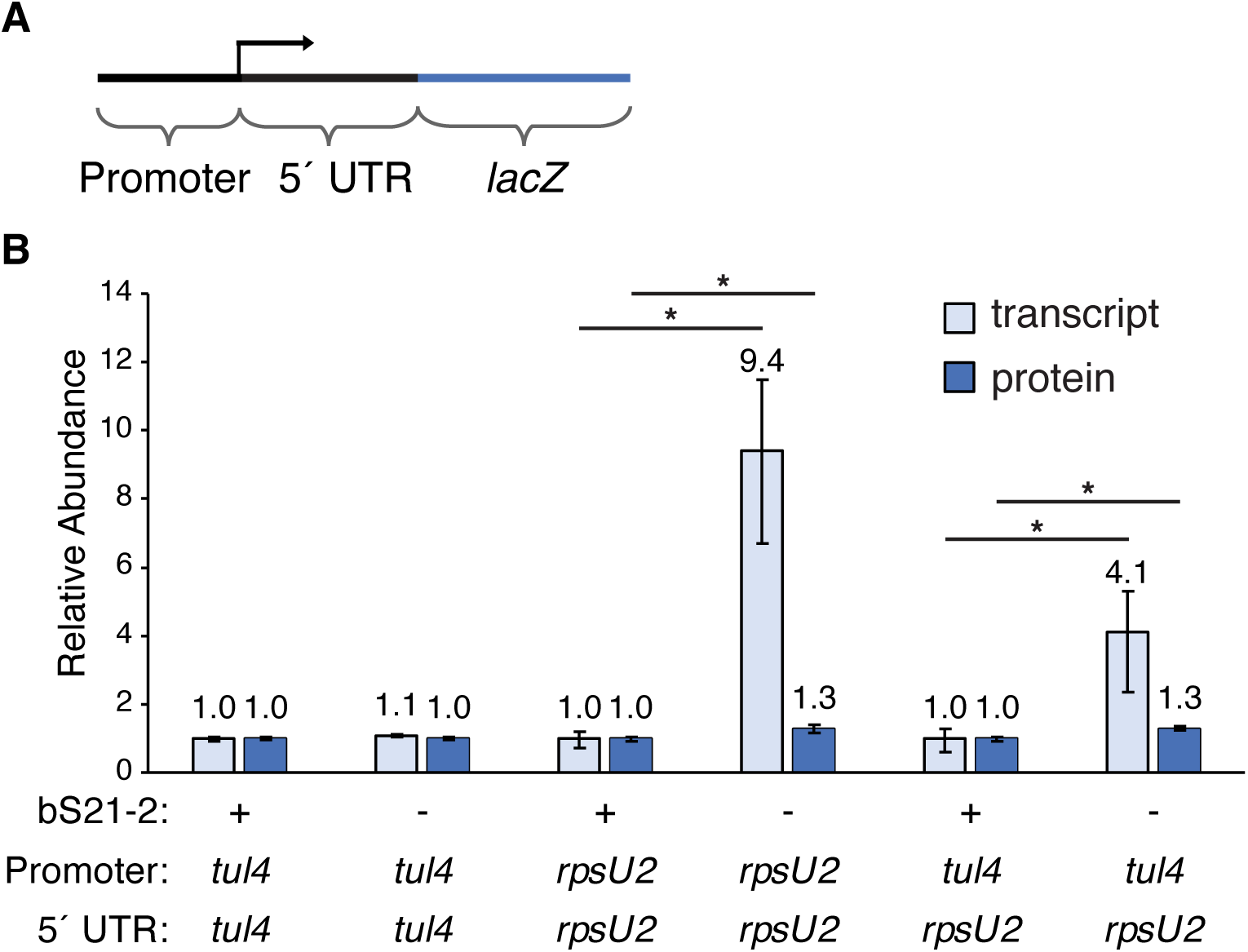
The *rpsU2* 5’ UTR mediates transcript abundance yet transcript and protein abundance changes are not correlated. (A) Diagram of translational reporters. (B) Relative mRNA and protein abundance for indicated products from translational fusions in cells with (+; wild-type) or without (-; Δ*rpsU2*) bS21-2. Quantitative RT-PCR was used to determine the rela-tive *lacZ* transcript normalized to the *tul4* gene. β-galactosidase activity was used to determine the relative protein abundance. Error bars represent 1 SD. Lines above bars indicate compari-sons between wild-type and Δ*rpsU2* of the same translational reporter, *p < 0.05. Experiments were repeated at least twice in biological triplicate and data from a representative experiment are shown.

There is evidence that this difference in relative transcription and translation is not unique to our reporter construct but reflects translation of the native *rpsU2* operon transcript. Specifically, while the abundance of the *rpsU2* operon transcript, including *yqeY*, is significantly increased in cells without bS21-2, proteomic analysis revealed no significant change in YqeY protein abundance (Trautmann and Ramsey, 2022). These findings suggest that while bS21-2 regulates the abundance of its own transcript, one or more additional factors control translation of the bS21-2 mRNA.

### The *rpsU2* 5’ UTR is sufficient for transcriptional autoregulation

There are a variety of mechanisms that control production of r-proteins and many of them depend on the mRNA leader sequence of the regulated r-protein. To determine if the 5’ UTR of *rpsU2* is sufficient to permit regulation by bS21-2, we created a translational fusion using the control promoter (*tul4*) driving expression of the 5’ UTR and first six codons of *rpsU2* in frame with *lacZ*. This reporter was similarly integrated onto the chromosome at the *att*Tn7 site in cells with and without bS21-2. We found that in cells without bS21-2, there was ∼4-fold more *lacZ* transcript compared to cells with bS21-2, suggesting that the *rpsU2* 5’ UTR is sufficient to allow control by bS21-2 (**Fig 3B**). Additionally, we observed the same disconnect between control of transcript abundance and translation. Despite the ∼4-fold increase in *lacZ* transcript abundance, we only found ∼30% more ß-galactosidase activity (**Fig 3B**). Thus, the *rpsU2* 5’ UTR is sufficient to lead to reduced mRNA abundance in cells with bS21-2 and regulation of translation by other factor(s).

### Stem-loops in the *rpsU2* mRNA do not influence regulation by bS21-2

We sought to investigate what features of the *rpsU2* 5’ UTR allow control by bS21-2. Secondary structures formed by mRNA leader sequences are often subject to regulatory control and the first 38 nt of the *rpsU2* 5’ UTR (−81 to −44) are predicted to form two stem-loop structures (**Fig 4A**). We considered the possibility that these stem-loops might be key for regulation of *rpsU2* mRNA abundance and generated additional translational reporters to test this hypothesis. Specifically, these reporters included either the full-length *rpsU2* 5’ UTR (81 nt) or truncated version (43 nt; “*rpsU2* Δloops”; **Fig 4B**) and the first six codons of *rpsU2* fused in-frame with *gfp*, driven by the control promoter (*tul4*). The reporters were integrated onto the chromosome at the *att*Tn7 site into cells with and without bS21-2 and we measured both *gfp* transcript abundance and GFP fluorescence (**Fig 4C**). Cells containing the GFP translational fusion with the full-length *rpsU2* 5’ UTR recapitulated our previous observations regarding regulation of the *rpsU2* mRNA: transcript abundance increases significantly in cells without bS21-2 (∼3-fold), but there is no similarly meaningful change in protein abundance. Notably, this outcome was unchanged in cells containing the truncated *rpsU2* 5’ UTR reporter construct (**Fig 4C**). These results indicate that the first 38 nt of the *rpsU2* mRNA, including the predicted stem-loops, are dispensable for regulation by bS21-2.

**Figure 4.**
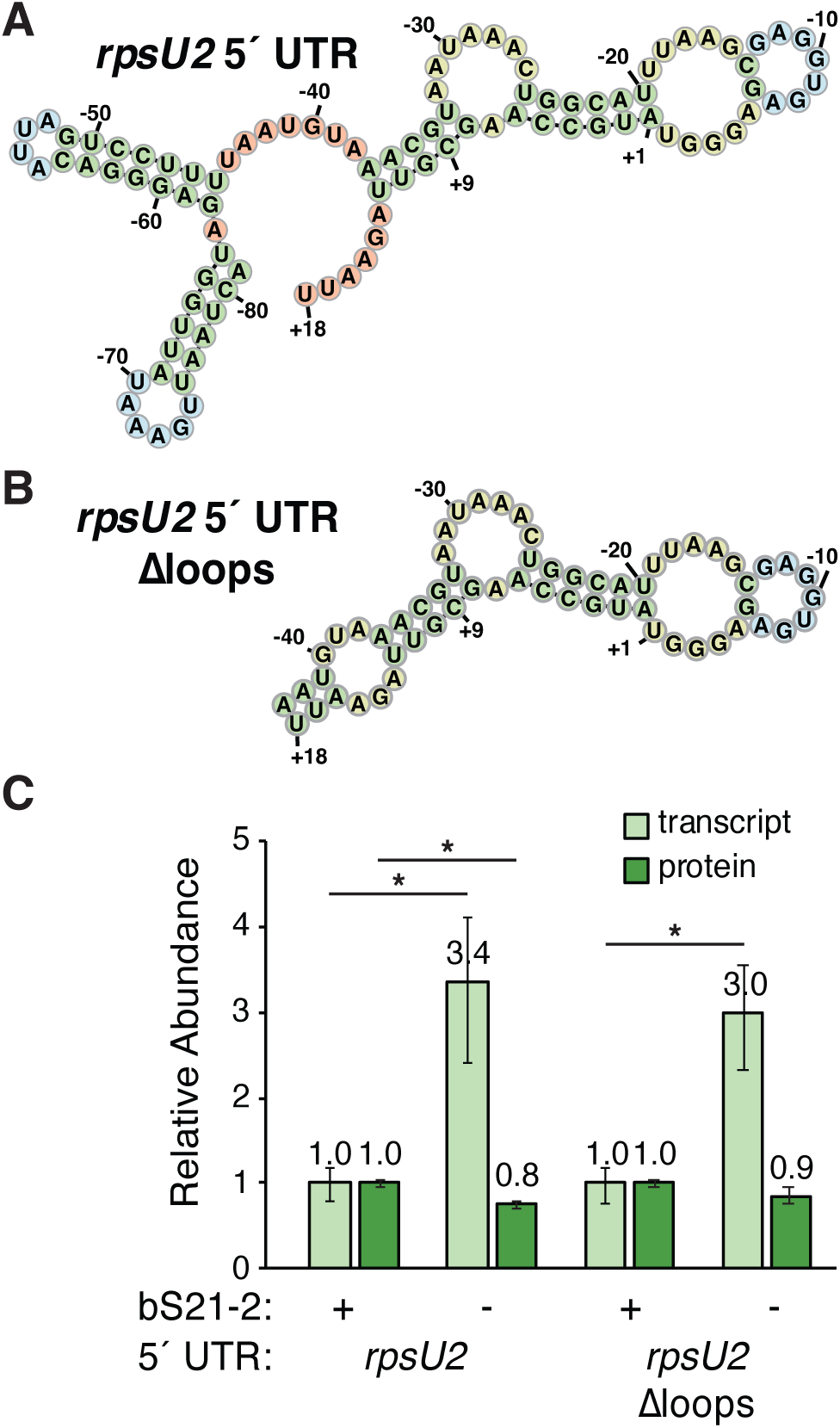
Predicted stem-loops in the *rpsU2* mRNA are dispensable for control by bS21-2. **(A)** Secondary structure predictions of wild-type and modified *rpsU2* 5 UTRs, generat-ed by MXfold2. **(B)** Relative mRNA and protein abundance for indicated products from transla-tional fusions in cells with (+; wild-type) or without (-; Δ*rpsU2*) bS21-2. Quantitative RT-PCR was used to determine the relative *gfp* transcript normalized to the *tul4* gene. Fluorescence was used to determine the relative protein abundance. Error bars represent 1 SD. Lines above bars indicate comparisons between wild-type and Δ*rpsU2* of the same translational reporter, *p < 0.05. Experiments were repeated at least twice in biological triplicate and data from a represen-tative experiment are shown.

### bS21-2 influences the stability of its own mRNA

The amount of any given transcript in a cell depends on two factors: the amount of new transcript produced and the rate of its degradation. Since the *rpsU2* 5’ UTR is sufficient for bS21-2-mediated changes in transcript abundance, we considered that stability, rather than changes in transcription rate, may be key in bS21-2-mediated regulation of *rpsU2* transcript abundance. Using a transcription inhibition assay, we compared the stability of the *rpsU2* operon transcript in cells with and without bS21-2 that harbor the *rpsU2*-*lacZ* translational reporter. Specifically, we added the RNA polymerase inhibitor rifampicin to mid-log phase cells to halt transcription and isolated RNA at multiple timepoints to assess the relative amount of *yqeY* transcript over time. This revealed that the *rpsU2* operon mRNA has a longer half-life in cells lacking bS21-2 (∼9 minutes compared to ∼1.5 minutes) and that bS21-2 decreases the stability of the *rpsU* mRNA, either directly or indirectly (**Fig. 5A**). Because the cells used in this experiment encode the *rpsU2*-*lacZ* reporter, we also tested the relative stability of the *lacZ* transcript and observed increased stability in cells lacking bS21-2 (**Fig. 5B**). From these observations, we conclude that the *rpsU2* leader sequence is sufficient to lead to increased mRNA stability in cells lacking bS21-2.

**Figure 5.**
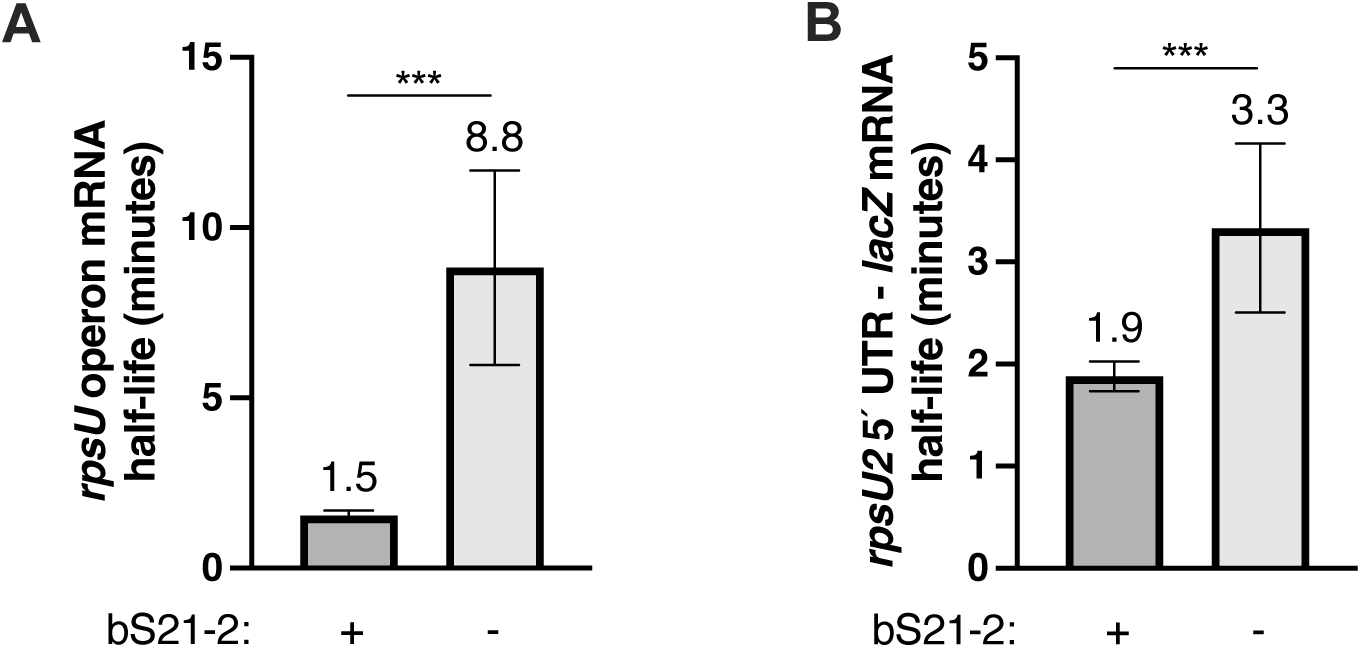
The presence of bS21-2 destabilizes transcripts with the *rpsU2* 5’ UTR. Half-lives of (A) *rpsU* operon mRNA as measured by *yqeY* transcript or (B) *lacZ* mRNA in cells containing P*_rpsU2_*-5’ UTR*_rpsU2_*-*lacZ* translational fusions with (+; wild-type) or without (-; Δ*rpsU2*) bS21-2. Linear regression analysis was used to calculate and compare half-life values. Error bars indicate standard error. Experiments were repeated at least twice in biological triplicate, and data from a representative experiment are shown. ***p<0.0001

## Discussion

In this study, we began to assess the factors that control production of different paralogs for the same ribosomal protein, bS21, in *F. tularensis*. We found that the presence of any of the three bS21 paralogs reduces the abundance of the bS21-2-encoding transcript, *rpsU2*, but neither bS21-1 nor bS21-3 influence the abundance of their own transcripts. Our work reveals that translation of the bS21-2 protein is further controlled by unknown factor(s), as large increases in *rpsU2* transcript in cells without bS21-2 does not lead to similar increases in protein abundance. We further determined that although the 5’ UTR of the *rpsU2* transcript is sufficient to lead to regulation by bS21-2, two predicted stem-loops in the leader sequence are not required. Finally, our work reveals that bS21-2 exerts at least some of its effects on *rpsU2* transcript abundance by altering mRNA stability. Our studies reveal that coordinated production of bS21 paralogs in *F. tularensis*, and particularly bS21-2, is complex and controlled at multiple steps.

In *F. tularensis*, all three bS21 paralogs participate in the regulation of the *rpsU2* operon transcript. This is revealed by the observation that cells without bS21-2 have a large increase in the transcript encoding bS21-2 and ectopic expression of any one of the three bS21 paralogs can reduce this steady-state transcript level to approximately wild-type levels. These bS21 paralogs are between 48-72% identical (Trautmann and Ramsey, 2022), and, given their similar influence on *rpsU2* mRNA, it raises the possibility that the conserved residues in bS21 paralogs are key for regulation of *rpsU2*, either directly or indirectly.

This first observation, that the presence of any of the three bS21 paralogs can reduce *rpsU2* transcript abundance, is consistent with well-described models of r-protein autogenous regulation (Meyer, 2018; Nomura et al., 1984). In these models, the amount of free r-protein is inversely correlated with its own production, such that as the r-protein becomes limiting for ribosome assembly, its transcript and/or translation increases. The autogenous r-protein regulators are generally primary binding proteins that bind specific rRNA structures with high affinity and bind a similar structure encoded within their leader sequence with lower affinity. They also often control the expression of multiple r-proteins encoded in the same operon, such that the imbalance of one r-protein regulates production of multiple r-proteins to restore stoichiometric balance between ribosomal components, as in uL1-mediated regulation of the uL1/uL11 operon in *E. coli* (Baughman and Nomura, 1983; Nevskaya et al., 2005).

In cells without bS21-2, the increase in *rpsU2* operon transcript is at least in part (and perhaps entirely) the result of increased mRNA stability. This raises the possibility that direct interaction between free bS21 and the *rpsU2* 5’ UTR decreases stability of the *rpsU2* operon transcript. This appears analogous to regulation of multiple r-proteins in *E. coli*, such as uS15. In this case, free uS15 leads to decreased stability of its own transcript, *rpsO*. The decrease in *rpsO* stability is due to uS15 stabilizing a pseudoknot structure in its leader sequence, precluding translation initiation (Philippe et al., 1993). The loss of mRNA protection by ribosomes allows ribonucleases increased access to the *rpsO* mRNA, resulting in higher degradation rates (Braun et al., 1998). Yet how bS21-2 leads to decreased stability of the *rpsU2* operon transcript is unclear. While it is possible that bS21 directly interacts with its own leader sequence, the structure of free bS21 is predicted to consist simply of two alpha helices connected by a short linker; there is no data to indicate if it can bind RNA alone. To date, there is no described bS21-binding motif in either the *rpsU2* leader or on the 16S rRNA and we have not identified elements in the *rpsU2* leader sequence that permits regulation by bS21 paralogs.

The outcome of the autogenous regulation model is a balance in r-protein abundance, with excess unincorporated r-protein inhibiting protein production while limited amounts of free r-protein trigger increased translation. Here, the comparison of bS21 and uS15 as regulators is less informative. While cells without bS21-2 have a large increase in *rpsU2* operon mRNA, there is no substantial change in the abundance of proteins encoded by this transcript. Thus, in the absence of bS21, we cannot attribute the increased stability of the *rpsU* mRNA to increased translation, because there is no apparent increase in translation, neither of the native operon or of translational fusions. We hypothesize that this discrepancy between relative transcript and protein abundance could be either (1) due to another factor (or factors) limiting translation initiation of mRNAs with the *rpsU2* leader sequence or (2) ribosomes lacking any bS21 paralog may be more abundant in cells without bS21-2 and these may specifically have reduced translation initiation efficiency on the *rpsU2* 5’ UTR.

While the 5’ UTR of the *rpsU2* transcript is sufficient for regulation by bS21-2, the specific elements of this mRNA that permit regulation have yet to be identified. In examining the contribution of two predicted stem-loops in the leader sequence, we narrowed down the sequence sufficient for regulation to 61 nt (18 nt encoding the first six codons and 43 nt of the leader sequence). Additional work will be required to identify what sequence and/or structural elements contribute to this regulation, as well as the molecular mechanism by which bS21-2 exerts its regulatory effects.

Production of ribosomes is energetically costly and the production of rRNA and r-proteins is tightly controlled by multiple inputs, ultimately linking ribosome number to growth rate. The bS21 paralogs in *F. tularensis* present an unusual but interesting case in r-protein production. It remains unclear why *Francisella* species maintain three different bS21 paralogs, nor what evolutionary processes led to their presence in the genome. It is unknown if the additional two paralogs were horizontally acquired or arose through duplication and divergence in a common ancestor. Regardless of their evolution, understanding how their production is coordinated with ribosome production and/or other processes in the cell, as well as the role of these paralogs in protein synthesis, provides a unique case study to understand how cells tightly control ribosome production and function.

## Materials and Methods

### Bacterial strains and growth conditions

*F. tularensis* subsp. *holarctica* LVS cells and derivatives were grown in Mueller-Hinton Broth supplemented with 0.025% iron pyrophosphate, 0.1% glucose, and 2% Isovitalex at 37°C shaking aerobically, or on cystine heart agar plates containing 1% hemoglobin (CHAH) at 37°C. *E. coli* XL1-Blue or PIR1 cells were grown in LB media or on LB plates. The antibiotics kanamycin, hygromycin, or nourseothricin were used for selection in *F. tularensis* LVS at 5 µg/mL or in *E. coli* at 50 µg/mL.

### Vector construction

A mini-Tn7 plasmid encoding the *rpsU2* promoter and 5’ UTR controlling *lacZ* was generated by modifying pKR68, a mini-Tn7 plasmid with a promoter-less *lacZ* gene (Trautman et al., 2023). Specifically, 258 bp of DNA upstream of the *rpsU2* coding sequence, including all of the intergenic sequence between the upstream, divergently-transcribed *dacB* and *rpsU2*, and the sequence specifying the first six codons of *rpsU2* was amplified from LVS genomic DNA using a 5’ primer specifying a KpnI site and a 3’ primer specifying a NotI site. This fragment was ligated into pKR68 digested with KpnI and NotI to generate pKR121. The mini-Tn7 plasmid pKR89, which replicates in *E. coli* using an R6K γ origin and encodes the *tul4* promoter and 5’ UTR was modified to create a reporter encoding the *tul4* promoter and *rpsU2* 5’ UTR (Trautmann et al., 2023). Specifically, a DNA sequence was synthesized by IDT specifying the 150 bp upstream of the *tul4* transcription start site followed by the DNA specifying the *rpsU2* 5’ UTR (81 bp upstream of the translation start site, as identified in Ramsey et al., 2015) and the first six codons. This fragment was amplified using a 5’ primer specifying a KpnI site and a 3’ primer specifying a NotI site, then ligated into pKR89 digested with KpnI and NotI to generate pKR123.

To avoid issues with overproduction of β-galactosidase in *E. coli* leading to toxicity, additional mini-Tn7 plasmids were created using sfGFP codon-optimized for *F. tularensis* LVS as a reporter, as in Trautmann et al., 2023. The DNA specifying the *tul4* promoter driving expression of the *rpsU2* 5’ UTR and first six codons was digested from pKR123 using KpnI and NotI, *gfp* was digested from pKR145 pF-*tul4* UTR-GFP (Trautmann et al., 2023) using NotI and BamHI, and the mini-Tn7 plasmid pKR121 was digested with KpnI and BamHI. The three fragments were ligated together to generate pKR184. To generate a version lacking the first 38 bp of the *rpsU2* 5’ UTR encoding two putative stem loops (Δloops), a primer was synthesized that specified the 3’ end of the *tul4* promoter, including the endogenous PacI site, followed by the sequence encoding the *rpsU2* 5’ UTR downstream of the putative loops. This primer, and one that would amplify part of *gfp* including the endogenously-encoded MfeI site, were used to amplify the *rpsU2* 5’ UTR lacking the putative loop region from pKR184. This fragment was ligated into pKR184 digested with PacI and MfeI to generate pKR191.

All plasmids used in this study are summarized in Table S1.

### Strains and strain construction

Derivatives of *F. tularensis* subsp. *holarctica* LVS used in this study are found in **Table 1**.

**Table 1.** *F. tularensis* LVS strains used in this study.

| Strain Number | Description | Genetic background | Integrated reporter plasmid |
| --- | --- | --- | --- |
| KRLVS111 | LVS $\Delta rpsU2$ Tn7::Ptul4-5'UTRtul4-lacZ aphA | LVS $\Delta rpsU2$ | pKR89 |
| KRLVS112 | LVS Tn7::Ptul4-5'UTRtul4-lacZ aphA | LVS | pKR89 |
| KRLVS148 | LVS $\Delta rpsU2$ Tn7::PrpsU2-5'UTRrpsU2-lacZ aphA | LVS $\Delta rpsU2$ | pKR121 |
| KRLVS149 | LVS Tn7::PrpsU2-5'UTRrpsU2-lacZ aphA | LVS | pKR121 |
| KRLVS150 | LVS $\Delta rpsU2$ Tn7::Ptul4-5'UTRrpsU2-lacZ aphA | LVS $\Delta rpsU2$ | pKR123 |
| KRLVS151 | LVS Tn7::Ptul4-5'UTRrpsU2-lacZ aphA | LVS | pKR123 |
| KRLVS279 | LVS Tn7::Ptul4-5'UTRrpsU2-gfp aphA | LVS | pKR184 |
| KRLVS280 | LVS $\Delta rpsU2$ Tn7::Ptul4-5'UTRrpsU2-gfp aphA | LVS $\Delta rpsU2$ | pKR184 |
| KRLVS281 | LVS Tn7::Ptul4-5'UTRrpsU2Δloops-gfp aphA | LVS | pKR191 |
| KRLVS282 | LVS $\Delta rpsU2$ Tn7::Ptul4-5'UTRrpsU2Δloops-gfp aphA | LVS $\Delta rpsU2$ | pKR191 |

Reporter constructs were integrated into the Tn7 site of *F. tularensis* LVS as previously described (LoVullo et al., 2009). Briefly, the Tn7 helper plasmid pMP720 was introduced by electroporation into *F. tularensis* cells with or without bS21 (LVS or LVS Δ*rpsU2*) and selection on CHAH plates containing hygromycin. Cells harboring the helper plasmid were subsequently electroporated with the appropriate mini-Tn7 plasmid and cells containing reporter constructs were selected by growth on CHAH plates with kanamycin. PCR was used to screen kanamycin-resistant colonies for plasmid integration at the attTn7 site and candidates were confirmed by amplification and sequencing of gDNA outside of the attTn7 site.

### mRNA stability assay

The stability of mRNAs was assessed essentially as described (Nguyen et al., 2020). Briefly, *F. tularensis* LVS cells containing the *rpsU2*-*lacZ* translational fusion with and without bS21-2 (KRLVS149 and KRLVS148) were grown to mid-log phase (OD_600_ = 0.3 – 0.4) in triplicate, and rifampicin was added to a final concentration of 50 µg/mL. After 0, 1, 2, 4, and 8 minutes, 7 mL cultures were snap frozen using liquid nitrogen and stored at −80°C until purification.

### RNA isolation, quantitative real-time PCR, and RNA-Seq

Cells of indicated *F. tularensis* LVS derivatives were grown to mid-log (OD_600_ = 0.3 – 0.4) in triplicate. Total nucleic acids were isolated using the Direct-zol RNA Kit (Zymo Research) following manufacturer’s protocols. Nucleic acids were treated with RQ1 DNase (Promega) and RNA was re-purified using the Direct-zol RNA Kit. cDNA was synthesized and quantitative real-time PCR was performed as described (Trautmann and Ramsey, 2022). Experiments were performed at least twice in biological triplicate. RNA, high-throughput sequencing, and data analysis for RNA-Seq was performed as described (Trautmann and Ramsey, 2022). RNA-Seq data are available in the National Center for Biotechnology Information Gene Expression Omnibus (NCBI GEO) under accession number GSE210766.

### β-Galactosidase assay

Indicated derivatives of *F. tularensis* LVS cells were grown to mid-log (OD_600_ = 0.3 – 0.4) in triplicate and ß-galactosidase activity was measured as previously described (Charity et al., 2009). Reactions were stopped after 120 minutes if no significant yellow color developed. Experiments were performed at least twice in biological triplicate.

### GFP assays

Indicated derivatives of *F. tularensis* LVS cells were grown to mid-log (OD_600_ = 0.3 – 0.4) and cells were pelleted and resuspended in PBS. Optical density (OD_600_) and fluorescence (excitation 495 nm, emission of 535 nm) were measured in technical triplicate using a SpectraMax® iD3 Multi-Mode Microplate Reader. Fluorescence readings were normalized to OD_600_ and the normalized fluorescent signal from cells containing an empty vector was subtracted. Experiments were conducted at least twice in biological triplicate.

## Acknowledgements

We thank Dr. Steven Gregory for his helpful comments on the manuscript and the other members of the Ramsey laboratory for support and helpful discussions. We also thank Janet Atoyan and the Rhode Island INBRE Molecular Informatics Core.

---

| Plasmid Name | Descriptive Name | Reference |
| --- | --- | --- |
| pF | pFNLTP6 | Maier et al., 2004 |
| pKR6 | pF-*rpsU1*-V | Trautmann et al, 2022 |
| pKR7 | pF-*rpsU2*-V | Trautmann et al, 2022 |
| pKR8 | pF-*rpsU3*-V | Trautmann et al, 2022 |
| pKR145 | pF-*tul4* UTR-GFP | Trautmann et al, 2023 |

Tn7 constructs
| Plasmid Name | Descriptive Name | Reference | Origin | Promoter | 5' UTR | Reporter | Backbone | Enzymes | Insert construction/PCR description |
| --- | --- | --- | --- | --- | --- | --- | --- | --- | --- |
| pMP720 | Tn7 helper plasmid | LoVullo et al., 2009 | pUC | - | - | - |  |  |  |
| pMP749 | Mini Tn7 plasmid | LoVullo et al., 2009 | pUC | - | - | - |  |  |  |
| pKR68 | Tn7 *lacZ* | Trautmann et al, 2023 | pUC | - | - | *lacZ* |  |  |  |
| pKR88 | Mini_Tn7_R6Kg | Trautmann et al, 2023 | R6K γ | - | - | *lacZ* |  |  |  |
| pKR89 | Tn7 *Ptul4*-5'UTR*tul4*-*lacZ* | Trautmann et al, 2023 | R6K γ | *tul4* | *tul4* | *lacZ* |  |  |  |
| pKR121 | Tn7 *PrpsU2*-5'UTR*rpsU2*-*lacZ* | This study | pUC | *rpsU2* | *rpsU2* | *lacZ* | pKR68 | KpnI/NotI | PCR of *rpsU2* promoter and 5'UTR from LVS gDNA |
| pKR123 | Tn7 *Ptul4*-5'UTR*rpsU2*-*lacZ* | This study | R6K γ | *tul4* | *rpsU2* | *lacZ* | pKR89 | KpnI/NotI | PCR of gBlock encoding *tul4* promoter fused to *rpsU2* 5' UTR |
| pKR184 | Tn7 *Ptul4*-5'UTR*rpsU2*-*gfp* | This study | pUC | *tul4* | *rpsU2* | *gfp* | pKR121 | BamHI/KpnI | *Ptul4*-5'UTR*rpsU2* fragment digested from pKR123 with KpnI/NotI; |
| pKR191 | Tn7 *Ptul4*-5'UTR*rpsU2* Δloops-*gfp* | This study | pUC | *tul4* | *rpsU2* Δloops | *gfp* | pKR184 | PacI/MfeI | Insert from pKR184 using KROL635 and KROL560 (PacI/MfeI digested) |

